# Template-based prediction of protein structure with deep learning

**DOI:** 10.1101/2020.06.02.129270

**Authors:** Haicang Zhang, Yufeng Shen

**Affiliations:** Department of Systems Biology, Columbia University, New York, NY, USA; Department of Biomedical Informatics, Columbia University, New York, NY, USA; JP Sulzberger Columbia Genome Center, Columbia University, New York, NY, USA; Program in Mathematical Genomics, Columbia University, New York, NY, USA

**Keywords:** protein structure prediction, protein threading, deep learning, deep residual neural network

## Abstract

Accurate prediction of protein structure is fundamentally important to understand biological function of proteins. Template-based modeling, including protein threading and homology modeling, is a popular method for protein tertiary structure prediction. However, accurate template-query alignment and template selection are still very challenging, especially for the proteins with only distant homologs available. We propose a new template-based modelling method called ThreaderAI to improve protein tertiary structure prediction. ThreaderAI formulates the task of aligning query sequence with template as the classical pixel classification problem in computer vision and naturally applies deep residual neural network in prediction. ThreaderAI first employs deep learning to predict residue-residue aligning probability matrix by integrating sequence profile, predicted sequential structural features, and predicted residueresidue contacts, and then builds template-query alignment by applying a dynamic programming algorithm on the probability matrix. We evaluated our methods both in generating accurate template-query alignment and protein threading. Experimental results show that ThreaderAI outperforms currently popular template-based modelling methods HHpred, CNFpred, and the latest contact-assisted method CEthreader, especially on the proteins that do not have close homologs with known structures. In particular, in terms of alignment accuracy measured with TM-score, ThreaderAI outperforms HHpred, CNFpred, and CEthreader by 56%, 13%, and 11%, respectively, on template-query pairs at the similarity of fold level from SCOPe data. And on CASP13’s TBM-hard data, ThreaderAI outperforms HHpred, CNFpred, and CEthreader by 16%, 9% and 8% in terms of TM-score, respectively. These results demonstrate that with the help of deep learning, ThreaderAI can significantly improve the accuracy of template-based structure prediction, especially for distant-homology proteins.

**Availability:** https://github.com/ShenLab/ThreaderAI

## 1 Introduction

Protein structure is fundamentally important to understand protein functions. Computational protein structure prediction remains one of the most challenging problems in structural bioinformatics. Recent progress in protein structure prediction showed that with the help of deep learning, it’s possible for free modelling (FM) methods to generate fold-level accuracy models of proteins lacking homologs in protein structure library^1–4^. Meanwhile, as both protein sequence and structure databases expand, template-based modelling (TBM) methods remain to be very popular and useful^5–7^ for the proteins with homologs available in protein structure library. TBM method predicts the structure of query protein by modifying the structural framework of its homologous protein with known structure in accordance with template-query alignment. The quality of TBM prediction critically relies on template-query alignment and template selection. It remains to be very challenging for TBM methods to predict structures accurately when only remote homologs which are conserved in structure but share low sequence similarity with query are available in structure library^5–7^.

The model accuracy of TBM method critically depends on protein features and the scoring functions that integrate these features. For protein features, sequence profiles, and protein secondary structures are widely used by exiting popular TBM methods such as HHpred^8^, CNFpred^9^, and Sparks-X^10^. As a result of recent progress in residue-residue contact prediction, contact information has been integrated by several recently developed methods such as DeepThreader^5^, CEthreader^6^, and EigenThreader^11^. For scoring functions, HHpred, Sparks-X, CEthreader, and several other methods used linear functions, while non-linear models such as Random Forest model in Boost-Threader^12^ and one-layer dense neural network in CNFpred have shown their advantages over linear models. Inspired by the success of non-linear models in TBM methods, we would like to study if we can improve TBM methods’ model accuracy using more advanced neural network architecture such as deep residual network which has proven very successful in protein residue-residue contacts prediction.

In this paper, we present a new method, called ThreaderAI, which uses a deep residual neural network to perform template-query alignment. More specifically, we formulate template-query alignment problem as the classical pixel classification problem in computation vision. We first adapt the deep residual neural network model to predict residual-residual aligning scoring matrix, and then we employ a dynamic programming algorithm on the predicted scoring matrix to generate the optimal template-query alignment.

## 2 Materials and Methods

### 2.1 Overview of the method

For a query protein, ThreaderAI predicts its tertiary structure through the following steps (Figure 1A). First, query protein is aligned to each template in the structure library using a deep residual neural network model and a dynamic programming algorithm. Second, all the alignments are ranked based on alignment scores. Third, the final tertiary structures of query are built using Modeller^13^ based on the top-ranked alignments.

**Figure 1.**
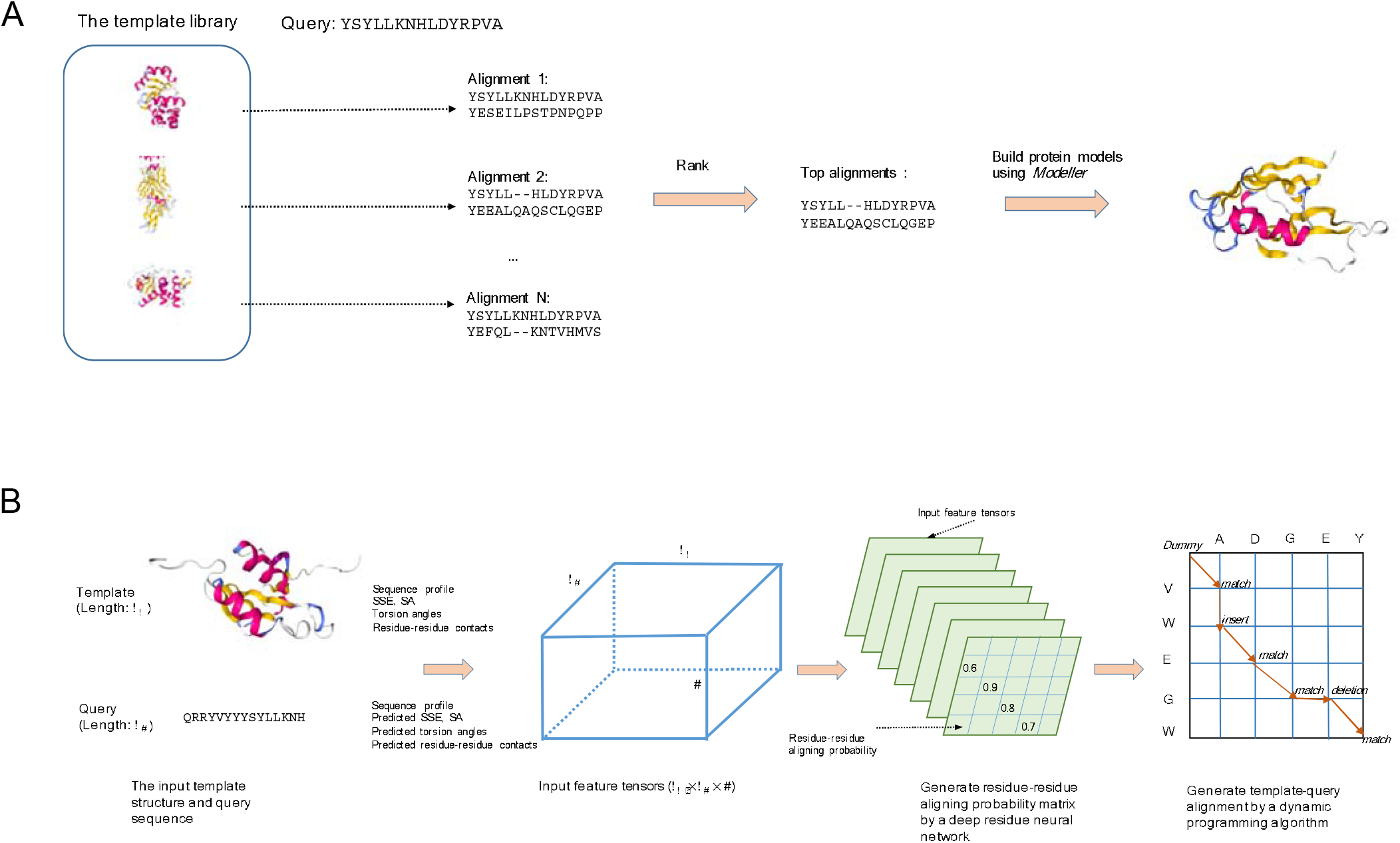
Overview of ThreaderAI. A. The procedure of protein structure prediction using ThreaderAI. B. The procedure of aligning query with template using a deep residual neural network model and a dynamic programming algorithm.

For TBM methods, the quality of query-template alignments critically determines the quality of predicted structures^5,9^. ThreaderAI uses a deep residue neural network model to generate template-sequence alignment (Figure 1B). First, protein features are extracted from both template and query. Second, a deep residue neural network model is used to generate residueresidue aligning probability matrix. Third, a dynamic programming algorithm is applied on the scoring matrix to generate the final template-query alignment.

### 2.2 Protein features

We included the following features as inputs for our deep residual neural network model (also see Table 1).

**Table 1.** Protein features used in ThreaderAI.

|  |  |
| --- | --- |
| sequence profiles (20×2 features) | Amino acid type distribution in multiple sequence alignment (20 features for template and 20 features for query) |
| sequential structural features only for template (13 features) | 8-class secondary structure types (8 features) |
|  | solvent accessibility (1 feature) |
|  | backbone dihedral angles (2 features) |
|  | contact numbers (2 features) |
| predicted sequential structural features for both template and query (8×2 features) | predicted 3-class secondary structure types (3 features) |
|  | predicted solvent accessibility (1 feature) |
|  | predicted backbone dihedral angles (2 features) |
|  | predicted residual level interfaces (1 feature) |
|  | predicted disordered regions (1 feature) |
| residual-residual contacts (8 features) | the dot products of the corresponding elements of top 8 eigenvectors of contact matrices of template and query |

*Sequence profile* (40 features): HHblits^14^ was used to generate the sequence profile for both template and query. The feature vector for each residue-residue aligning pair is from the concatenation of the sequence profiles of template and query.

*Sequential Structural features* (29 features): For template, we generated its 8-class secondary structure types, real-valued solvent accessibility, and backbone dihedral angles using DSSP^15^. We also calculated the contact numbers of template with *C_α_-C_α_* and *C_β_-C_β_* distances of 8A□ as threshold. And for Glycine, we only used its *C_α_* coordinates. For query, we predicted its 3-class secondary structure types, real-valued solvent accessibility, backbone dihedral angles, disordered regions, and residual level interfaces using NetSurfP2^16^. We also predicted these sequential structural features for template. The features of a residue-residue aligning pair are the concatenation of the structural properties of these two residues.

*Residue-residue contacts* (8 features): The residue-residue contacts of template are defined as the residue pairs with *C_β_-C_β_* distance less than 8A□. For query, we predict its contact map using ResPRE^17^. The eigenvectors and eigenvalues of the residue-residue contact matrix can capture the intrinsic properties of protein’s tertiary structure and have been used as features by recently developed threading methods^6,11^. Given contact matrix *M* of the protein, the *i*th residue can be represented as 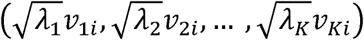 where *λ_j_* and *V_j_* is the *j*th eigenvalue and eigenvector of matrix *M,* respectively. Here we set *K* as 8. Given template and query’s contact matrices *M_T_* and *M_Q_*, the features of the *i*th residue of template aligning the *j*th residue of query are defined as 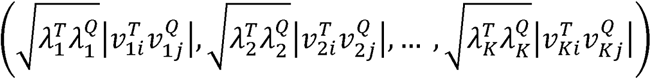. Heuristically, we set the sign of each eigenvector as positive. Previous methods^6,11^ enumerated a total of 2^*K*^ possible alignments to decide the sign of each involved eigenvector which is very time-consuming and infeasible for neural network-based models.

### 2.3 Neural network architecture

We employed a deep residual neural network^18^ (ResNet) model to predict residue-residue aligning probability matrix. ResNet has proven very successful in computer vision and also in structural bioinformatics. First, convolutional layers in ResNet are capable of extracting hierarchical features or spatial patterns from images or image-like data automatically. Second, the residual component in ResNet can efficiently mitigate the issue of vanishing/exploding gradients and makes it possible to train an ultra-deep neural network model on a large scale of training data.

Specifically, for a template-query pair, the input feature tensor for our neural network model has dimensions of *L_T_* x *L_Q_* x *d* where *L_T_* and *L_Q_* denotes the lengths of template and query, respectively, and *d* is the number of features for each residue-residue pair. And the output for our model has dimensions of *L_T_* x *L_Q_* each element of which representing residue-residue aligning probability. Our model includes 16 residue blocks^18^ each of which includes 2 convolutional layers. Each convolutional layers used 16 filters and a kernel size of 3 × 3. We used ELU^19^ as nonlinear activation function. Sigmoid function was used as the final layer to output residueresidue aligning probabilitites.

### 2.4 Alignment labels and training loss function

We built training template-query pairs from proteins with known structures. For each template-query pair in training data, we used DeepAlign^20^ to generate its structural alignments as ground truth. For a template with a length of *L_T_* and a query with a length of *L_Q_*, there are *L_T_* X *L_Q_* residue-residue pairs in total, in which the aligned pairs in the structural alignment are labeled as positives while the others as negatives.

Instead of using binary labels directly, we weighted^9^ the conservation of aligned residue pairs using local TM-score^21^. Given a structure alignment of two proteins and the corresponding superimposition, the local TM-score of an aligned residue pair *T_i_* and *Q_j_* is defined as follows:

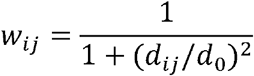

where *d_ij_* is the distance deviation between the two aligned residues and *d*_0_ is a normalization constant depending only on protein length. The TM-score ranges from 0 to 1, with higher values indicating more highly conserved aligned positions. And for a gap in the alignment, the local TM-score *w_ij_* is equal to 0.

The labels from the structure alignments are highly imbalanced in which the ratio of negatives over positives is proportional to the lengths of template and query. To mitigate this imbalanced labeling issue, we weighted the aligned pairs in the reference alignments with the average length of template and query.

We used cross-entropy loss as our training loss function which is defined as follows:

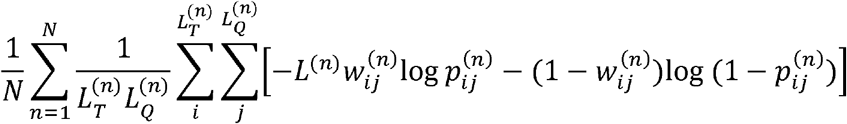

where *N* is the number of protein pairs in training data and *n* iterates over all training samples, and *L*^(*n*)^ equals 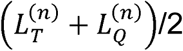 meaning the average length of template and query, and 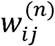 and 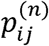 are residue-residue aligning probability from our neural network model and local TM-score, respectively.

### 2.5 Training algorithm

We used AdamW algorithm^22^ to minimize the objective function with a weight decay rate of 1e-4. For the warmup stage, we increased the learning rate from 0 to 0.01 over the first 2 epochs. We also decayed the learning rate to 1e-4 with a polynomial decay policy in the following 16 epochs^22^. Early-stopping with validation error as a metric was performed during training. The model architecture and training algorithm was implemented by TensorFlow2^23^ and run on 3 NVIDIA GeForce-1080 GPUs in parallel. We set training batch size as 2 and we didn’t try a larger batch size due to the limited GPU memory.

### 2.6 Maximum accuracy algorithm

Given the residue-residue aligning probability matrix of *L_T_* × *L_Q_* from our neural network model, we used a dynamic programming algorithm called Maximum Accuracy algorithm (MAC)^8,14^ to generate the final template-query alignment. MAC creates the local alignment through maximizing the sum of probabilities for each residue pair to be aligned minus a penalty *α* which can control the alignment greediness. To find the best MAC alignment path, an optimal subalignment score matrix *S* is calculated recursively using the probability *p_ij_* as substitution scores:

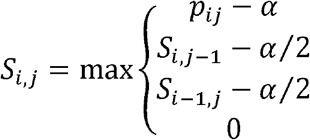

Then standard traceback procedure of dynamic programming^24^ was then applied on the score matrix *S* to generate the optimal local alignment. We rank the template-query alignments based on the optimal alignment scores from MAC.

### 2.7 Dealing with proteins of variable lengths

Our model has an architecture of fully convolutional neural network^25^ in which no fully-connected layers were used. As a result, the number of parameters of our model is independent of the lengths of both template and query. Hence, our model can deal with proteins of variable lengths. In particular, zero paddings were applied so that each training sample in the same minibatch has the same size. We also filtered out the padded positions when we aggregated the final training loss.

### 2.8 Training and test data

We built the training set, validation set, and independent testing set from proteins in SCOPe40. We also included CASP13 data for testing.

#### 2.8.1 Training data

We prepared template-query pairs from SCOPe40^26^. First, for testing purpose, we excluded the domains which share larger than 25% sequence identity with the domains in CASP13 data^7^. Here we used MMseqs2^27^ to evaluate sequence identity with the default E-value of 1e-3. Second, we excluded families with single domains. Third, for each class of *α, β, α/β*, and *α + β* of SCOPe40, we randomly selected 5 folds as independent testing data and the left folds as training data. The testing and training template-query pairs were generated from testing and training folds respectively.

Template-query pairs at the similarity of fold, superfamily and family levels were generated separately. When generating family level pairs, at most 10 pairs were randomly selected for each family. And when generating superfamily and fold level pairs, for each family pairs from the same fold, we randomly selected 1 domain from each family as its representative to form pairs. And all protein pairs with TM-score less than 0.3 were excluded. Finally, we have 53734 training pairs and 2000 validation pairs from the training folds, and 3106 pairs from testing folds.

#### 2.8.2 Test data

We used two test datasets to test ThreaderAI in terms of alignment accuracy and protein threading performance, respectively. For testing alignment accuracy, we used 3106 template-query pairs (denoted as SCOPe3K data) created together with training pairs and validation pairs (see section 2.8.1). The testing template-query pairs belong to different folds with training and validation pairs. The second test set consists of 61 officially-defined CASP13^7^ target domains under the category of Template-Based Modelling (TBM). The CPSP13 TBM data are divided into two groups by difficulty level: TBM-easy (40 targets) and TBM-hard (21 targets). We used

To test the threading performance of ThreaderAI using CASP13 TBM data, we built our template database from PDB90 in which any two proteins share less than 90% sequence identity. We only included the structures deposited before CASP13. We also excluded the structures with more than 800 amino acids and the structures with more than 50% unobserved residues. Finally, our template library includes 50099 proteins.

### 2.9 Evaluating metrics

#### 2.9.1 Evaluating alignment accuracy

For a query protein and one of a candidate template from the template library, we evaluated the alignment accuracy by evaluating the quality of the structure built from this alignment. In particular, for each template-query pair, we first used ThreaderAI to generate an alignment, then built a 3D structure for the query using MODELLER^13^ based on the alignment, and finally evaluate the similarity between the predicted structure and the ground truth structure. Here, we evaluated the quality of a 3D model by GDT^28^ and TM-score, two widely used metric for measuring the similarity of two protein structures. GDT score is calculated based on the largest set of residue-residue pairs falling in a defined distance cutoff when superposing these two structures. GDT ranges from 0 to 100, but we normalize it by 100 so that it has a scale between 0 and 1. TM-score is designed to be length-independent by introducing a length-dependent normalization factor. TM-score ranges from 0 to 1 with 1 indicating the perfect model quality.

#### 2.9.2 Evaluating threading performance

We evaluated threading performance by measuring the quality of 3D models built from the topranked templates. Specifically, for a query protein, we used ThreaderAI to generate alignments for all the templates in template library, ranked these alignments by alignment scores and then built 3D models using MODELLER from the top five alignments. Finally, we evaluated the quality of the first-ranked and the best of top five 3D models by TM-score and GDT.

### 2.10 Compare with previously published methods

We compared ThreaderAI with several widely used threading methods including HHpred^8^, CNFpred^9^, and CEthreader^6^, a new threading method built upon contacts predicted by ResPre^17^. Here, HHpred was run with the option mact 0.1, real secondary structures for template, and predicted secondary structures for query proteins. And CEthreader was run with the mode of EigenProfileAlign in which sequence profile, secondary structures, and contact maps are used. For protein threading, we used CEthreader’s suggested strategy to speedup. That is, we first run CEthreader’s greedy algorithm and then selected top the 1000 templates for refinement using its enumerative algorithm. DeepThreader^5^ is another recently developed threading software in which a linear function was used to combine local potentials from CNFpred and pairwise potentials from predicted residue-residue contacts. DeepThreader’s performance wasn’t shown here because its package is unavailable to the public. To be fair, for all methods we used the same template database (see section 2.8.2) and used HHblits^14^ to build sequence profiles against sequence database uniclust30_2017_10 built before CASP13. We used HHblits’ utility script to convert HHBlits’ profile format to BLAST’s^29^ profile format used by CNFpred.

## 3 Results

### 3.1 Alignment accuracy on SCOPe3K data

Based on SCOPe’s hierarchical classification for proteins, we split all the template-query pairs into three groups: the pairs similar at family level, at superfamily level, and at fold level. Two proteins are similar at fold level if both query and template belong to the same fold but different super families. The similarity at superfamily level and family level are defined in the same way. Two proteins similar at fold level are conserved in structure but diverges in sequence, and are usually considered as remote homologs, while two protein similar at family level share high sequence similarity and are usually considered as close homologs.

As shown in Table 2 and Figure 2, on SCOPe3K data, ThreaderAI outperforms all other competitors including HHpred, CNFpred, and CEThreader in terms of alignment accuracy. In particular, ThreaderAI achieved average TM-score and GDT of 0.510 and 0.437, respectively. In terms of TM-score, ThreaderAI outperforms HHpred, CNFpred, and CEthreader by 23%, 9%, and 12%, respectively. The advantage of ThreaderAI over the second-best method is the largest when the similarity between template and query falls into fold level, which indicates ThreaderAI’s power in modelling of remote homologs. In particular, at the fold level, ThreaderAI outperforms HHpred, CNFpred, and CEthreader by 56%, 13%, and 11% in terms of TM-score, respectively. The advantages of ThreaderAI over other methods decreases at the family level, which is not surprising since it is easy to align two closely-related proteins. At the superfamily level, ThreaderAI outperforms HHpred, CNFpred, and CEthreader by 18%, 9%, and 14% in terms of TM-score, respectively.

**Figure 2.**
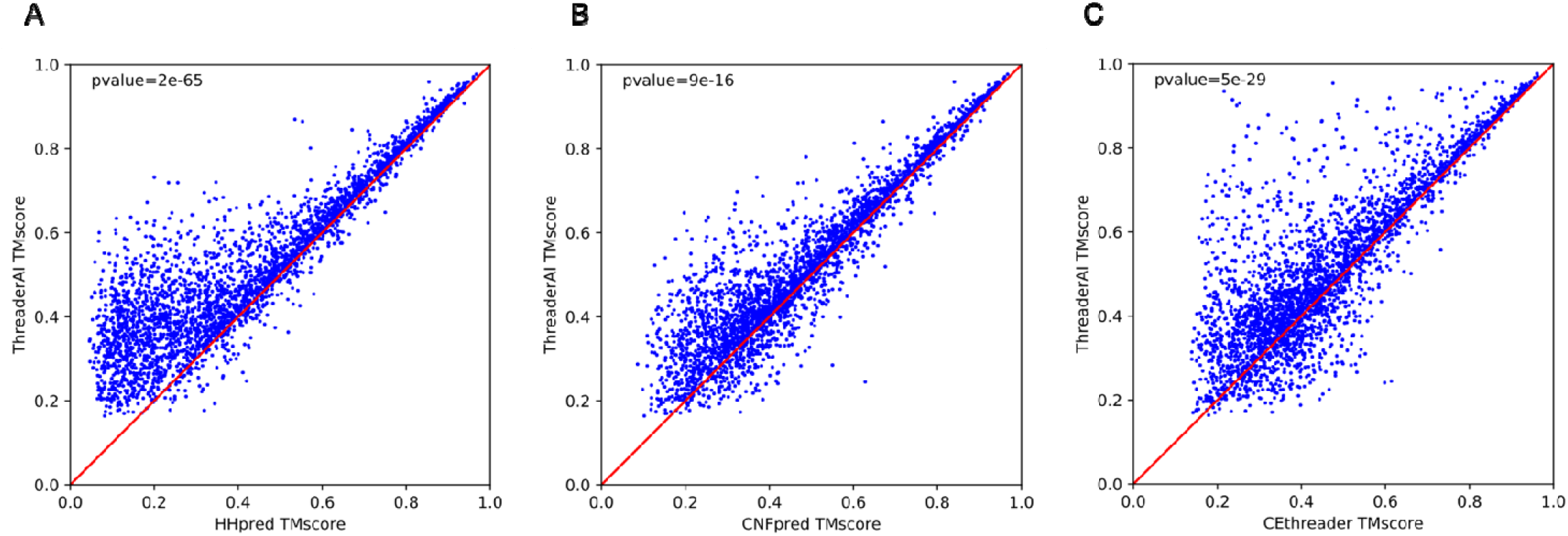
Comparison of ThreaderAI and previously published methods using alignment accuracy on SCOPe3K. Each point in the figure represents alignment accuracy of ThreaderAI versus the other competing method.

**Table 2.** Alignment accuracy measured by TM-score and GDT on SCOPe3K data

|  | Fold level |  | Superfamily level |  | Family level |  | All |  |
| --- | --- | --- | --- | --- | --- | --- | --- | --- |
|  | TM-score | GDT | TM-score | GDT | TM-score | GDT | TM-score | GDT |
| ThreaderAI | 0.419 | 0.348 | 0.483 | 0.409 | 0.705 | 0.633 | 0.510 | 0.437 |
| HHpred | 0.268 | 0.226 | 0.411 | 0.353 | 0.675 | 0.607 | 0.416 | 0.362 |
| CNFpred | 0.371 | 0.307 | 0.443 | 0.375 | 0.681 | 0.612 | 0.470 | 0.404 |
| CEthreader | 0.377 | 0.313 | 0.425 | 0.356 | 0.641 | 0.568 | 0.456 | 0.389 |

We also used a t-test to assess the statistical significance of the comparison results. On 3206 template-query pairs, in terms of TM-score, the *p*-values between ThreaderAI and HHpred, CNFpred, and CEthreader are 2e-65, 9e-16, and 5e-29, respectively. Figure 2 shows more details on the difference of alignment accuracy between ThreaderAI and the competing methods. In terms of TM-score, ThreaderAI achieved better alignment quality than CNFpred for 2743, 2395, and 2343 pairs, while worse for 363, 711, and 763 pairs, respectively. It confirms that ThreaderAI can generate better alignments than our competing methods.

### 3.2 Threading performance on CASP13 data

We further evaluated the threading performance of our method on the 61 CASP13 TBM domains. Among the TBM domains, 40 and 21 domains belong to the categories of TBM-easy and TMB-hard, respectively. Here ThreaderAI and all competitors used the same template database (see section 2.8.2).

As shown in Table 3, on all TBM targets, ThreaderAI outperforms all the competing methods no matter whether the models are built from the first-ranked or the best of top five templates. ThreaderAI achieves a TM-score 0.761 for first-ranked models, which outperforms HHpred, CNFpred, and CEthreader 10%, 5%, and 6%, respectively. Overall, ThreaderAI shows larger advantages on the TBM-hard group in which only remote homologs are available. Specifically, on TBM-hard group, ThreaderAI outperforms HHpred, CNFpred, and CEthreader by 16%, 9%, and 8%, respectively. This again indicates ThreaderAI’s great advantages in modelling of remote homologs.

**Table 3.** Threading performance on 61 CASP13 TBM domains. Each cell shows the average quality of the 3D models built from the first-ranked and the best of top five templates.

|  | TBM-easy |  | TBM-hard |  | TBM-All |  |
| --- | --- | --- | --- | --- | --- | --- |
|  | TM-score | GDT | TM-score | GDT | TM-score | GDT |
| ThreaderAI | 0.813/0.831 | 0.752/0.776 | 0.663/0.702 | 0.554/0.608 | 0.761/0.787 | 0.684/0.719 |
| HHpred | 0.753/0.779 | 0.704/0.733 | 0.570/0.629 | 0.477/0.554 | 0.690/0.728 | 0.626/0.672 |
| CNFpred | 0.785/0.805 | 0.727/0.751 | 0.611/0.659 | 0.520/0.571 | 0.726/0.755 | 0.656/0.689 |
| CEthreader | 0.770/0.795 | 0.715/0.740 | 0.615/0.665 | 0.533/0.582 | 0.717/0.751 | 0.653/0.686 |

### 3.3 Running time

With the help of GPUs’ computational power, ThreaderAI is very efficient in protein threading (Figure 3). As far as we know, ThreaderAI is the first template-based modelling method which can take advantage of GPUs. ThreaderAI first uses 3 GeForce-1080 GPUs to generate the scoring matrices for all templates in the template library and meanwhile uses 4 CPU cores to maintain the data stream for the model. And then ThreaderAI runs the Maximum Accuracy Algorithm for all scoring matrices on 1 CPU core.

**Figure 3.**
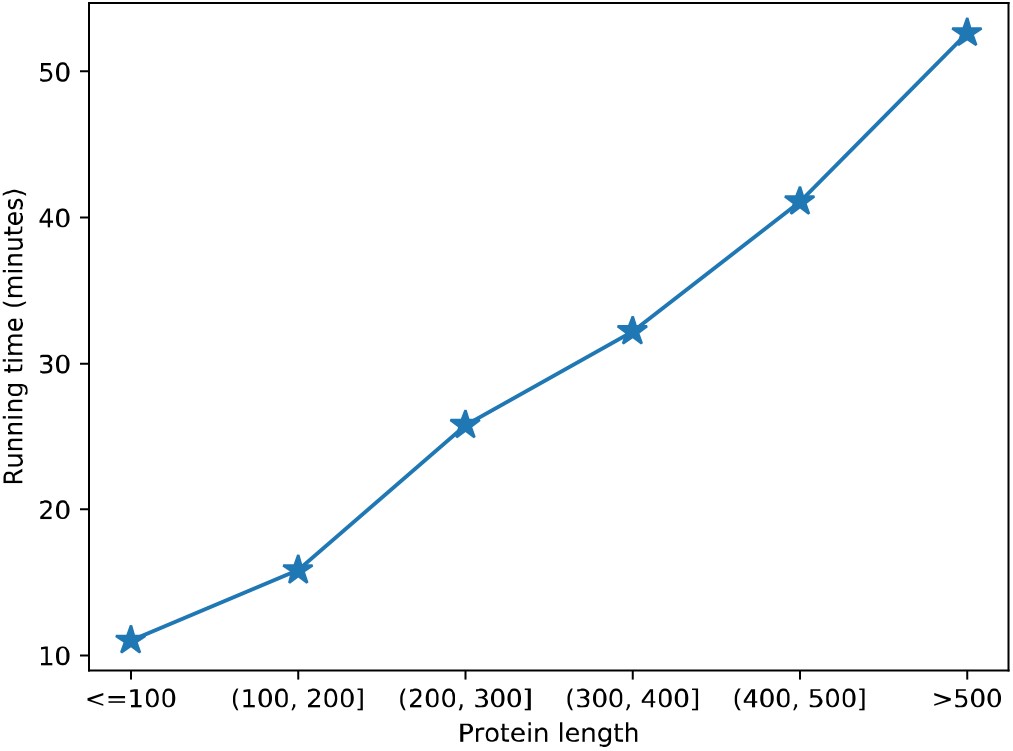
The running time of ThreaderAI searching query protein in CASP13 data against PDB90. Here we split the data into several groups based on protein lengths. Y-axis is the mean running time in minutes for each group.

The running time of ThreaderAI mainly depends on protein length. The protein threading can be finished within 20 minutes for proteins with less than 200 amino acids. And it takes ThreaderAI less than 1 hour to finish protein threading even for the proteins with length larger than 500. ThreaderAI is highly scalable as it can use more GPUs.

## 4 Discussion

We developed ThreaderAI, a new template-based method for predicting protein structure using a deep residual neural network. We show that Threader outperforms the existing popular TBM methods including HHpred, CNFpred, and CEthreader, in both alignment accuracy and threading performance, especially on proteins that only have remote homologs with known structure. In particular, ThrederAI outperforms CNFpred, another neural network based-method, in which only one dense layer is used. This demonstrates that advanced neural network models are more capable of capturing complex sequence-structure relationship.

ThreaderAI formulates the template-query alignment problem as the classical pixel classification problem in computer vision. To fulfill this, residue-residue pair scoring is separated from alignment generation. It’s still possible to design an end-to-end model to produce template-query alignment by combining a deep residual neural network and a chain graphical model such as Hidden Markov Model^30^ and Condition Random Fields^31^. However, in the hybrid model, the gradients of neural network will entangle with the gradients of chain graphical model which makes it very inefficient to train a deep model on a large scale of training samples^32^.

ThreaderAI could be improved in several directions. First, besides deep residual neural network, other deep learning models such as deep autoregressive models^33^ may improve alignment accuracy. Second, deep attention model^34^ may provide a more efficient way to integrate residueresidue contact information. ThreaderAI integrates residue-residue contacts indirectly by including the eigenvectors of the contact matrix in which the sign of eigenvectors are decided very heuristically. Local potentials and pairwise potentials related to the residue-residue contact pairs and non-contacting pairs can be weighted directly with the help of attention mechanisms.

